# Structural basis of the interaction between ESV1 and LESV from *Arabidopsis thaliana* with starch glucans

**DOI:** 10.1101/2023.06.09.544376

**Authors:** Rayan Osman, Mélanie Bossu, David Dauvillée, Corentin Spriet, Chun Liu, Samuel Zeeman, Christophe D’Hulst, Coralie Bompard

## Abstract

Starch is the major energy storage compound in plants. Whether it is transient or stored, it is accumulated in the form of insoluble, semi-crystalline granules. The structure of these granules is related to the structure of the main component: amylopectin. Amylopectin consists of linear polymers of glucose units linked by α-1,4 bonds, forming double helices that combine to form the semi-crystalline lamellae of the granules, and α-1,6 branching points that form the amorphous lamellae. This particular structure of amylopectin is linked to the action of isoamylases, which cut the excess of branching points and allow the granules to be structured. For a long time, it was thought that the action of these enzymes was responsible for the structuring of starch granules. Recently, two new proteins, LESV and ESV1, have been characterized and are involved in the phase transition of amylopectin (LESV) or in the maintenance of the granule structure (ESV1). These proteins share a tryptophan-rich domain folded into an antiparallel β-sheet that is particularly well suited to bind amylopectin double helices. In this paper we present the structural study of these interactions using integrative structural biology approaches and show that LESV, in contrast to ESV1 can intervenes during amylopectin biosynthesis.

## Introduction

In plants, starch is a storage polymer made of glucose residues. It accumulates as water-insoluble, partly crystalline granules which size varies from 0.1 to 100 μm in diameter. In seeds, roots and tubers, storage starch serves as a long-term carbohydrate reserve, used to fuel germination or seasonal re-growth. In leaves, transitory starch accumulates in chloroplasts of photosynthetic cells during the day and is used as carbon and energy source during the night. This process is under a photoperiodic control guaranteeing progressive consumption of the starch reserve until dawn (Stitt and Zeeman 2012). Starch granules are made up of two polymers of glucose residues, namely amylose and amylopectin that adopt different 3D structures (for review (Pfister and Zeeman 2016)) and have different physicochemical properties. Amylopectin accounts for 70-80% of the starch content, with the remainder being amylose. Amylopectin, is organized as α-1,4-glucans linked one to another by α-1,6-bonds (= branching points), while amylose is poorly branched (<1% α-1,6 bonds)(Pfister and Zeeman 2016; Pérez and Bertoft 2010).

Amylopectin is synthesized by the concerted activities of soluble starch synthases (SSs), starch branching enzymes (BEs) and starch debranching enzymes (DBEs). Soluble starch synthases (SSs) transfer the glucose residue of ADP-Glucose (the precursor molecule for starch biosynthesis in plants) to the non-reducing end of an elongating glucan (Xie et al. 2018). Branch points are introduced by BEs which cleave an α-1,4 bond of a pre-existing glucan and transfer the malto-oligosaccharide located toward the non-reducing end on a neighboring glucan (intermolecular reaction) or to the other part of the cleaved glucan (intramolecular reaction) by an α-1,6 bond (Sawada et al. 2014). Debranching enzymes comprise isoamylases (Isa) and pullulanases. Specific Isa isoforms are involved in the synthesis of amylopectin by hydrolyzing α-1,6 bonds of a developing glycogen-like nascent amylopectin molecule in amorphous lamellae to ordinate the branch points and allow the amylopectin to crystallize (Ball et al. 1996; Myers et al. 2000; Wattebled et al. 2005).

Within amylopectin, the entwining of adjacent chains folded into double helices gives rise to both secondary and tertiary structures of the molecules. These structures align and pack into dense, crystalline lamellae, which alternate with amorphous lamellae that contain the branch points and chains connecting the crystalline layers within the starch granule. The resulting regular pattern of crystalline (∼6 nm) and amorphous (∼3 nm) layers is a common feature of plant starches, and is believed to underlie the frequently observed 9-10 nm repeat structure (Buleon et al. 1998).

Although the main steps of starch metabolism have been quite well described to date, the processes defining starch granule size and shape, and the internal matrix organization remain to be elucidated (Pfister and Zeeman 2016). It has been widely assumed that the starch granule matrix formation involves self-organization physical process events during the early stages of granule formation (Waight et al. 2000; Ziegler, Creek, and Runt 2005). However, two recently discovered non-catalytic proteins, Early Starvation 1 (ESV1) and Like Early Starvation 1 (LESV), have been described to be involved respectively in starch granules stabilization and structuration of crystalline layers of starch (Liu, Pfister, Osman, Ritter, Heutinck, Sharma, Eicke, Fischer-Stettler, et al. 2023). ESV1 and LESV were discovered in *Arabidopsis thaliana* leaves and potato tuber (Feike et al. 2016; Helle et al. 2018). The wide occurrence and conservation of these two proteins in different plants indicates that they have a fundamental function in starch metabolism although they don’t have an identified enzymatic activity. The implication of ESV1 and LESV in starch metabolism has been demonstrated after the analysis of KO mutant lines of *Arabidopsis* in which the starch phenotype has been specifically altered. In addition, ESV1 and LESV appear to have an opposite function since *Arabidopsis* lines overexpressing LESV have a phenotype similar to that of *esv1* KO mutant (Feike et al. 2016). The presence and/or level of these proteins are crucial for normal starch biogenesis and degradation in plants and this process varies between plant organs. Zeeman’s team proposed that *At*ESV1 and *At*LESV could be involved in determining the conformation of the starch granule matrix structure (Feike et al. 2016).

Protein sequence analysis fails to identify already known catalytic domains or any other conserved domain (as Carbohydrate binding modules, CBM). Sequence analysis of ESV1 and LESV shows that both proteins have an analogous domain of about 240 amino acids located at the C-terminus of both proteins. However, their N-terminal regions, on the other hand, are of different length (about 130 amino-acids in ESV1 and 304 in LESV including the transit peptide) and do not share sequence homology. The C-terminal domains contain numerous Tryptophan and aromatic amino acid residues organized in conserved repeated motifs also containing acidic residues (D,E). The presence of these repeated motifs could constitute binding sites for numerous glucans or mediate interaction with long glucans as starch components (Feike et al. 2016). In a very recent work, the function of both proteins was further investigated by a combination of structural and functional studies(Liu, Pfister, Osman, Ritter, Heutinck, Sharma, Eicke, Fischer-Stettler, et al. 2023). The structures of ESV1 and LESV were modeled using Alphafold (Jumper et al. 2021) and completed by biophysical approaches (Liu, Pfister, Osman, Ritter, Heutinck, Sharma, Eicke, Fischer-Stettler, et al. 2023). The result showed an unique and common fold for the conserved C-terminal domain (containing about 250 amino acid residues) of both proteins. The tryptophan-rich regions of both *At*ESV1 and *At*LESV proteins are predicted with high confidence, folding into an extended planar β-sheet consisting of 16 twisted antiparallel β- strands. Localizing the conserved motifs of aromatic and acidic residues within these predicted structures revealed them as aligning into linear stripes regularly spaced and running across both sides of the β-sheet, perpendicular to the β-strands. The distance between aromatic stripes is about 14 Å and 70 Å long which is globally the size of a double helix of amylopectin in the crystalline phase of starch granules (Buleon et al. 1998). It has been proposed that this domain could constitute a new carbohydrate binding surface capable of binding at least two double helices of amylopectin molecule on each side of the β-sheet. Synthetic biology approaches in yeast and *in-vivo* experiments in *Arabidopsis*, provide direct evidence that LESV is directly involved in the phase transition of amylopectin. To do that the described carbohydrate binding surface would allow LESV to bind several linear chains of amylopectin and thus promote the organization of the crystal phase of starch granules. This domain would allow ESV1 to maintain the organization of glucans in neo-formed granules and to limit their enzymatic degradation during the daytime phase (Liu, Pfister, Osman, Ritter, Heutinck, Sharma, Eicke, Fischer-Stettler, et al. 2023).

In this work, we analyzed the interaction of LESV and ESV1 with starch glucans by a combination of biochemical and biophysical approaches and describe the effect of this interaction on 3D structures of both proteins.

## Methods

### Cloning, expression and purification of proteins

AtLESV and AtESV1 have been cloned, expressed and purified as described previously (Liu, Pfister, Osman, Ritter, Heutinck, Sharma, Eicke, Fischer-Stettler, et al. 2023). Briefly the gene of both proteins have been synthesized with codons optimized for production in *E. coli* and inserted in plasmids allowing the insertion of a 6His-tag. For both proteins the N-terminal transit peptides (56 and 95 residues for AtLESV and ESV1 respectively) have been truncated. Both proteins have been expressed in *Escherichia coli*. Purification of the proteins has been performed by a first step of Immobilized Metal Affinity Chromatography followed by a second purification step through size exclusion chromatography using a HiLoad 16/60 Superdex 200 (Cytiva) column pre-equilibrated with 50 mM Tris pH 8, 150 mM NaCl, 10% glycerol, 2 mM DTT (w/v) for LESV or a dialysis step against 50 mM Tris pH 7.5, 100 mM NaCl, 10% [v/v] glycerol, 2 mM DTT for ESV1. The monodispersity of the obtained protein solution has been assessed by dynamic light scattering (DLS) using a zetasizer pro (Malvern Pananlytical). For structural study, protein samples were concentrated using Vivaspin centrifugal concentrator with a 10 kDa cut-off (sartorius). Protein concentrations were determined using a Nanodrop Spectrophotometer (ND1000) from Thermo Scientific.

### Glucan solutions preparation

For EMSA experiments, amylose 1% (w/v), amylopectin 1% (w/v) and glycogen 1% (w/v) stock solutions used for these experiments were prepared as follows. 0.1g of amylose (from potato, Sigma) was dissolved in 1 mL of 2M sodium hydroxide (NaOH) and vortexed to ensure complete solubilization. After addition of 2 ml of deionized water, the solution was neutralized by 1 ml of 2M hydrochloric acid (HCl). The final volume was then adjusted to 10 ml with water and the solution was heated 5 min at 50°C and vortexed until complete dissolution. Amylopectin (100 mg, from potato, Sigma) was resuspended in 10 mL of distilled water and then subjected to autoclaving to produce a homogeneous solution. One hundred mg of glycogen (from oyster, Sigma) powder was dissolved in 10 mL of distilled water and then stirred at 40°C until full homogenization. For CD experiments, glucan stock solutions were prepared with the same protocol by replacing water with protein buffer.

### ElectroMobility Shift Assays (EMSA)

ESV1 and LESV (1 μg) and a reference protein (1μg) (ref uniprot Q7W019 with no affinity for glucans) were loaded on 8% polyacrylamide gels containing increasing concentrations of glucan solutions (from 0.1 to 0.3% for amylopectin and amylose and 0.1 to 0.5% for glycogen) and submitted to an electrophoresis in native conditions at 4°C at 15V cm^-1^ for 2h in 25mM Tris, 192 mM Glycine, pH8.3 migration buffer. Gels were stained with InstantBlue^TM^ (Expedeon) after rinsing in deionized water. The affinity of LESV and ESV1 for the different glucans present at different concentrations in the gels has been estimated by the migration shift of these proteins compared to the stable migration of the control protein which has no affinity for the glucans.

### Synchrotron radiation circular dichroism

Synchrotron radiation circular dichroism (SR-CD) spectra were measured at the DISCO beamline of the SOLEIL Synchrotron (Gif-sur-Yvette, France). 5 µL of ESV1 at 6.1 mg/ml and 2 µl of AtLESV at 13.4 mg/ml were deposited between 2 CaF_2_ coverslips with a pathlength of 20 µm and 10 µm respectively (Refregiers et al. 2012). The beam size of 4 × 4 mm and the photon-flux per nm step of 2 × 10^10^ photons s^−1^ in the spectral band from 270– 170 nm prevented radiation-induced damage (Miles et al. 2008). CD spectra were acquired using IGOR software (WaveMetrics). Protein and buffer spectra were collected consecutively and are the mean of 3 accumulations. The buffer baseline was recorded sequentially and subtracted from the spectra before taking into account the concentration in residues. Before measurements the molar elliptical extinction coefficient of Ammonium *d*-10- Camphorsulfonate (CSA) has been measured on the beamline and used as standard for calibration of all data measurements (Miles, Wien, and Wallace 2004). Data processing was conducted using CDToolX software (Miles and Wallace 2018). The influence of the different glucans on the structure of LESV and ESV1 was studied by incubating the protein/glucan mixtures for 2h and measuring the spectra under the same conditions as the native proteins. The mixtures were made with 80% protein solution and 20% of amylose or amylopectin solutions 1% w/v. 5 µL of ESV1 at 6.1 mg/ml/glucans mixtures and 2 µl of AtLESV at 13.4 mg/ml/glucan mixtures were deposited between 2 CaF_2_ coverslips with a pathlength of 20 µm and 10 µm respectively (Refregiers et al. 2012). Spectra containing 80% of protein buffer and 20% glucan solutions were subtracted from the protein/glucan spectra before CSA calibration.

Temperature scans were realized to check the protein stabilization by glucan interaction. CD spectra were collected from 20-30 to 90°C and processed as described above with 3-5°C temperature increases and 3 min of equilibration time.

The secondary structure element content of each protein alone or in the presence of glucans was estimated using BestSel (Micsonai et al. 2015).

### Molecular Modelling

Protein structures of LESV and ESV1 have been modelled using AlphaFold2 (Jumper et al. 2021). For each protein, five different models have been computed and ranked by global pLDDT. The five molecular models generated were superposed and used to evaluate the possible position of dynamic regions. Molecular models with best pLDDT have been used for figures and further molecular docking.

### Small-angle-X-ray-Scattering (SAXS)

Protein sample solutions were centrifuged for 10 min at 10,000 g prior to X-ray analysis to remove aggregates. SAXS experiments were conducted on the SWING beamline at Synchrotron SOLEIL (λ= 1.033 Å). All solutions were mixed in a fixed-temperature (15°C) quartz capillary. The monodisperse sample solutions of proteins were injected onto a size exclusion column (David and Perez 2009) (SEC-3, 150 Å; Agilent) using an Agilent HPLC system and eluted into the capillary cell at a flow rate of 0.3 ml min^-1^. Then 50 µl of protein samples were injected for SAXS measurements. A large number of frames were collected during the first minutes of the elution and were averaged to account for buffer scattering. The latter was subtracted from selected frames corresponding to the main elution peak. Data reduction to absolute units, frame averaging, and subtraction were done using FOXTROT (David and Perez 2009). All subsequent data processing, analysis, and modeling steps were carried out using programs of the ATSAS suite (Franke et al. 2017). The radius of gyration (*Rg*) was derived by the Guinier approximation using PRIMUS (Franke et al. 2017). The program GNOM (Svergun 1992) was used to compute the pair-distance distribution functions [*P*(*r*)] and feature the maximum dimension of the macromolecule (*D*max). DAMMIF/DAMAVER/DAMFILT (Volkov and Svergun 2003) were used to model protein envelopes and structure. BUNCH (Petoukhov and Svergun 2005) was used to model the missing parts of the proteins that were not assigned by Alphafold (Jumper et al. 2021).

### Docking

Alphafold model structure of the conserved C-terminal domain of ESV1 and LESV was used as the target in order to model the binding position of the model of double helix of amylopectin obtained from Polysac3DB (CERMAV https://polysac3db.cermav.cnrs.fr). A generic algorithm was used for the search step within a sphere of 10 Å centred on the tryptophane stripes of the conserved β-sheet. The scoring function was based on the ChemPLP forcefield, as used by GOLD (Jones et al. 1997). All the parameters were kept by default. A subsequent energy minimization was performed on the best model using the Amber forcefield. All figures representing molecular structures of proteins and ligands were generated using Pymol (The PyMOL Molecular Graphics System, version 1.8.0.0 Schrödinger, LLC).

### 3D imaging

For this experiment we used *waxy* maize starch granules (Roquette, France). *waxy* starch was obtained from plants lacking granule-bound starch synthase (GBSS;(Tsai 1974)), the enzyme that catalyzes the biosynthesis of amylose and whose presence in the granules is responsible of their autofluorescence. Starch granules were washed with water, acetone and ethanol in order to remove any phenol contaminants coming from plastic storage as described in (Tawil et al. 2011). ESV1 and LESV binding was followed by UV microscopy on starch granules and followed by both visible and UV microscopy with the same protocol described in (Jamme et al. 2014). Excitation was set-up at λ=280 nm with an emission filter at 329-351 nm (FF01-340/22, Semorock) to visualize the tryptophan emission (at 345 nm) of bound proteins. The acquisition time was 10 seconds for each emission fluorescence image and 0.2 s for visible images. For 3D purpose, Z slices (along the optical axis) were recorded with a step size of 300 nm over 40 µm Z range under µManager control (Edelstein et al. 2010). Using imaging analysis software (Huygens, SVI, NL), deconvolution images were calculated by PSF deconvolution treatment. Images were coupled to classical light imaging of the starch granule morphology. Images were analyzed using FiJi (Schindelin et al. 2012). Noise was removed from acquired 3D stacks using a median 1 filter. A substack of “in focus” images was then selected and summed together. To compensate for field inhomogeneity, a FFT bandbass filter was then applied.

## Results

### The N-ter domain of LESV contains helices with folding and position depending on protein environment

To go beyond the Alphafold models available for LESV and ESV1 on the Alphafold platform, we recalculated a set of five models for both proteins using Alphafold 2. Specifically, we calculated in-house the templates for the N-terminally truncated constructs made for the expression of the two proteins used in the CD and SAXS experiments (Liu, Pfister, Osman, Ritter, Heutinck, Sharma, Eicke, Fischer-Stettler, et al. 2023). The predicted regions with a high degree of confidence (pLDDT > 90) for ESV1 (amino acids from 142 to 395) and LESV (amino acids from 318 to 573) remain identical in the 5 models generated and consist mainly of the β-stranded tryptophan-rich C-terminal region described in a recent work. This result suggests a high stability for this domain, which is predicted to be a new carbohydrate-binding domain (Liu, Pfister, Osman, Ritter, Heutinck, Sharma, Eicke, Fischer-Stettler, et al. 2023). In these 5 models we were interested in the predicted structures for the less conserved domains. As previously described, the regions close to the C-terminal β-sheet of ESV1 (46 amino acid residues at the N-terminus and the polyproline region at the C-terminus), which are poorly conserved, are predicted to be disordered. In the C-terminal region of the β-sheet, located on one of its faces (Face A), an α-helical region (residues 378 to 395) is predicted with a high degree of confidence in its ÷structure and position relative to the β-sheet. This helix is also present in LESV (residues 555 to 578) with the same degree of confidence. Both regions contain 4 conserved amino acids (Liu, Pfister, Osman, Ritter, Heutinck, Sharma, Eicke, Fischer-Stettler, et al. 2023).

The structure of the N-terminal region of LESV is predicted with low to very low confidence. However, it has been shown that three helical regions located in an island of conservation are predicted with pLDDT >70 (Liu, Pfister, Osman, Ritter, Heutinck, Sharma, Eicke, Fischer-Stettler, et al. 2023). Only one of these forms a long helix (residues 245 to 273) whose position relative to the β-sheet, is predicted with high confidence (Figure 1A). This helix is located on the Face A of the β-sheet. It is predicted with the same confidence in all models generated by Alphafold, but its position may vary from one model to another, suggesting that it may be modified according to the protein’s environment. For other helices, although predicted with pLDDT > 70, their positions are not equivalent (position and limits) between the five models calculated here. This result may indicate that the N-terminal domain of LESV is disordered and susceptible to induce helix folding under certain conditions.

**Figure 1:**
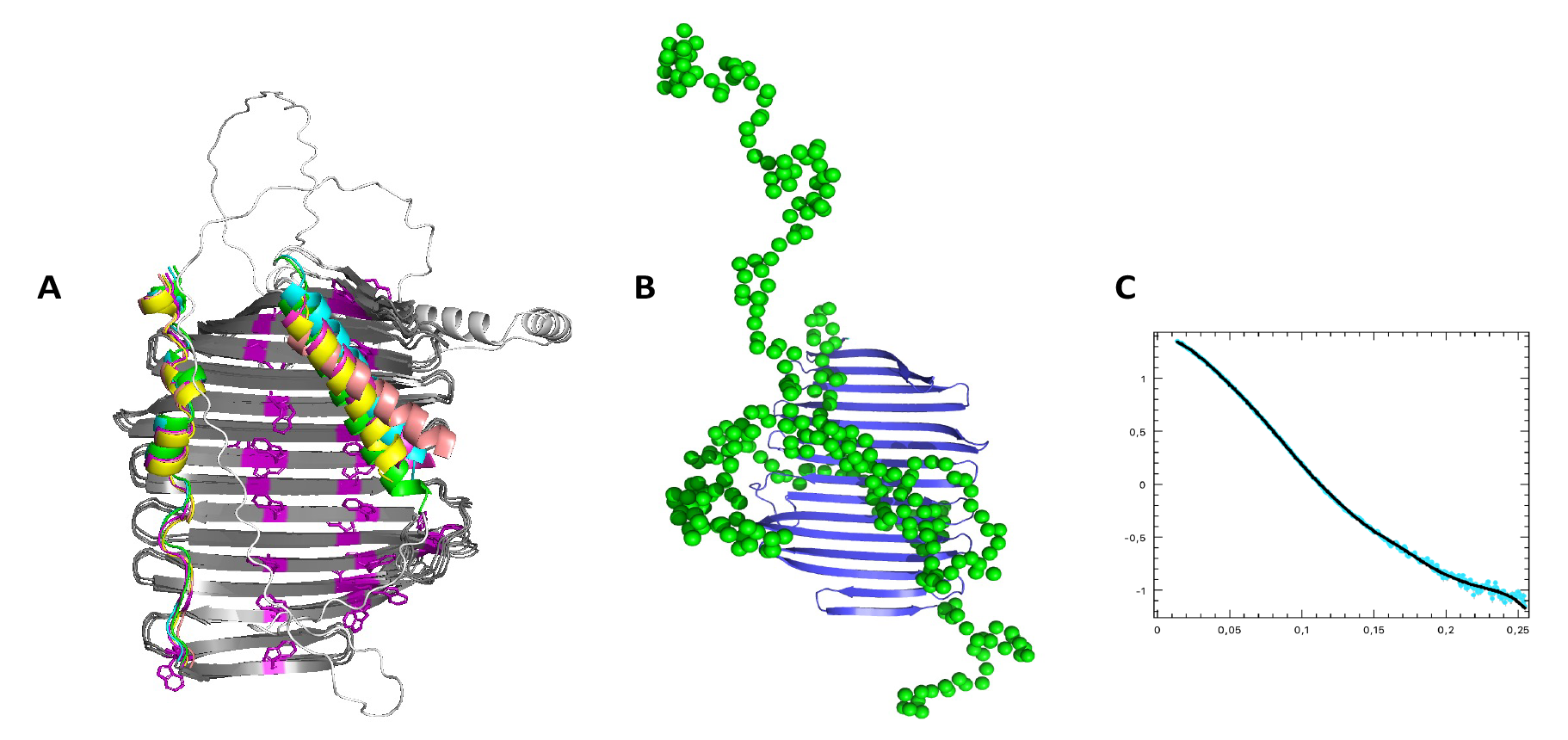
*Structure dynamics of LESV from Arabidopsis thaliana.* The structure is represented in cartoon. A) Superposition of the 5 molecular models of AtLESV calculated with Alphafold 2. Only regions with pLDDT > 70 are shown except for the first model (light gray). Regions common to all 5 models with pLDDT > 90 are colored dark grey. The helices of the 5 models are colored in light grey, salmon, cyan, yellow and green. The common helix with ESV1 is on the left, the long helix specific for LESV is on the right. B) *ab initio* model of the N-terminal domain of LESV computed from SAXS data using the software BUNCH. The conserved C-terminal domain is represented as cartoon and colored in dark blue, the N- terminal domain model is represented as spheres (One sphere by amino acid residue) and colored in green C) Superposition of SAXS experimental data obtained for LESV (cyan) and calculated curve from BUNCH model (black) with ξ^2^= 3.8Å.

We have analyzed the structure of the Face A of the β-sheet of ESV1 in the region equivalent to that occupied by helix 245-273 on LESV. Alphafold does not predict a long helix at this location, but a loop (amino acids 109 to 138) with a pLDDT> 70 and an expected positional error between < 3 Å (Figure S1). The helix and loop are stabilized by numerous interactions with the amino acids of the β-sheet of both proteins, including the conserved aromatic and acidic residues that cover half the height of the sheet. These two structures are located on the same side of the β-sheet as the C-terminal helices conserved in both proteins and described above. Their presence in this configuration is not compatible with the binding of the amylopectin double helices, resulting in a polarity in the glucan-binding domain: one side is accessible, the other is not.

### The N-terminal domain of LESV is partially disordered and folds close to the tryptophan rich domain

In order to obtain more information on the folding and structural state of the two proteins in solution, we continued the analysis of the SAXS data presented in (Liu, Pfister, Osman, Ritter, Heutinck, Sharma, Eicke, Fischer-Stettler, et al. 2023). In this previous study the envelopes of both ESV1 and LESV were revealed with SAXS analysis. For the ESV1 construct, which contains virtually only the conserved tryptophan-rich domain, we showed that the Alphafold model for this domain is consistent with the calculated molecular envelope of the protein. This was also the case for LESV, but the presence of the N-terminal domain, for which Alphafold predicts no reliable structure except for a few conserved helices, makes comparison difficult (Liu, Pfister, Osman, Ritter, Heutinck, Sharma, Eicke, Fischer-Stettler, et al. 2023).

In order to visualize and localize the N-terminal domain of LESV, we performed an *ab initio* modelling based on the structure of the high-resolution C-terminal domain model given by Alphafold and the SAXS data using BUNCH, a program that allows to model missing parts of a protein from SAXS data (Petoukhov and Svergun 2005). The result obtained for LESV is shown in **Figure 1B**. The obtained model, which fits the SAXS data with a high degree of confidence (χ^2^= 3.8) (**Figure 1C or supplementary),** shows that the end of the C- and N-terminal domain (between 50 and 80 residues) are rather disordered and emerge from the overall structure, while the rest of the domain is organized around or close to the β-sheet in structures that could be compatible with the helices predicted by Alphafold. The presence of this domain around, or at least close to, the β-sheet is likely to affect glucan-binding and could explain the differences in function between the two proteins described in (Liu, Pfister, Osman, Ritter, Heutinck, Sharma, Eicke, Fischer-Stettler, et al. 2023).

### ESV1 and LESV interact differently with α-1,4-linked glucose polymers

To better understand the specificities of ESV1 and LESV and their interaction with starch glucans, we performed several experiments. The structural heterogeneity of amylose and amylopectin solutions as well as their high viscosity precluded their use in conventional structural biology approaches (SAXS or X-ray crystallography) or SPR.

Alternatively, the interaction of amylose and amylopectin with ESV1 and LESV was investigated using EMSA. EMSA is a rapid and sensitive method to detect protein-glucan interactions. It is based on the observation that the electrophoretic mobility of a protein in acrylamide gels can be modified in gels containing increasing concentrations of a ligand. Protein interaction with the ligand present in the gel will lead to a decrease in its electrophoretic mobility and the corresponding band will be shifted to higher molecular weight. The intensity of this shift will be proportional to the concentration of glucan in the gel. First, we followed the influence of increasing concentrations of amylose or amylopectin on the electrophoretic mobility of LESV and ESV1.

The results are shown in **Figure 2**. On native polyacrylamide gels containing amylopectin, both ESV1 and LESV show a large shift for the ESV1 and LESV bands which increases with amylopectin concentration (Figure 2A). As low as 0.1% final concentration of amylopectin into the gel was enough to induce a strong reduction in electrophoretic mobility compared to the control. ESV1 and LESV were similarly delayed and the relevant bands were located just below the stacking gel demonstrating a strong affinity of the two proteins for amylopectin.

**Figure 2:**
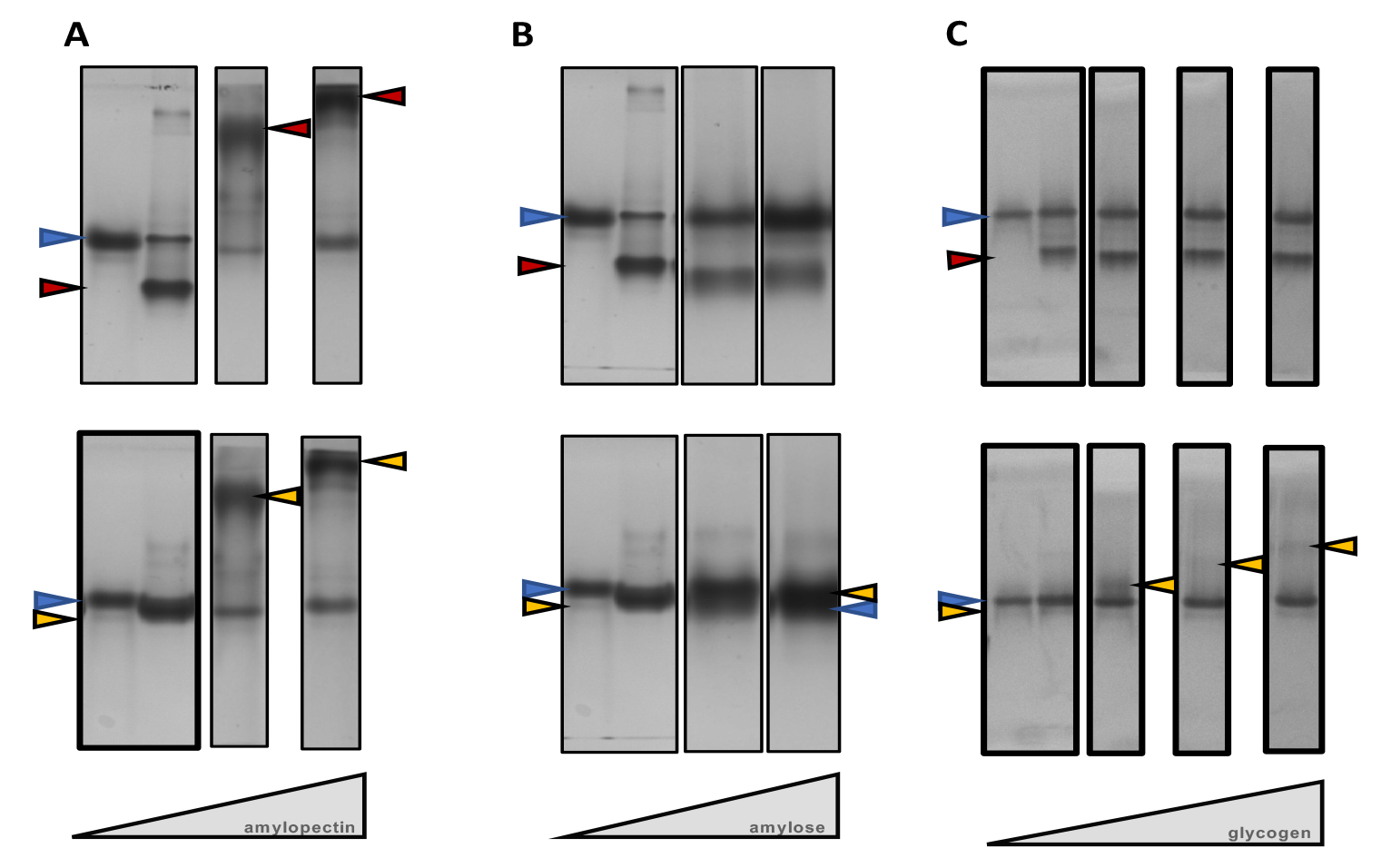
*EMSA gels analyzing the interaction between ESV1 (top) or LESV (bottom)* and A) amylopectin B) amylose, or C) glycogen. Blue, red and yellow arrows indicate the bands corresponding to the reference protein, ESV1 and LESV respectively.

In contrast, no electrophoretic mobility differences were found for LESV and ESV1 in native gels containing amylose in the same range of concentration. The few differences that could be seen were not concentration dependent (Figure 2B). This result suggests that LESV and ESV1 have no affinity for amylose under the tested conditions.

Amylose has longer chains than amylopectin and much fewer branching points. To try to understand the reasons for the differences in affinity between the two starch components, we decided to test the affinity of LESV and ESV1 for glycogen (α-1,4-linked glucans highly branched by α-1-6-linkages). The results obtained for glycogen are shown in **Figure 2C**. Three different final concentrations of glycogen (0.1%, 0.3%, 0.5%) were added to 8% acrylamide native gels. A shift of the LESV protein band is clearly observed in the gel containing 0.1% glycogen compared to the reference protein band. This shift is accentuated when the glycogen concentration is increased, suggesting here again that LESV has some kind of affinity for this branched glucan. ESV1, on the other hand, does not show any change in its electrophoretic mobility in the presence of glycogen, eliminating a potential affinity between ESV1 and glycogen.

### Binding of amylopectin to LESV causes α-helices to appear in the protein structure

In order to better characterize the mode of interaction of ESV1 and LESV with glucans, and in particular to highlight conformational changes of the proteins’ structure during complex formation, we performed an SR-CD study. CD is the method of choice for analyzing the structure of a protein by visualizing its content of secondary structural elements. It also allows the study of interactions between proteins and their ligands, especially when these latter lead to structural changes. SR-CD extends the limits of typical CD spectroscopy by providing an extended spectral range and improving signal-to-noise ratio and faster data acquisition, even in the presence of absorbing elements such as buffers and salts (Hussain, Javorti, and Siligardi 2012; Hussain, Longo, and Siligardi 2018)

Quantitative analysis of CD spectra also makes it possible to predict the secondary structure content of a protein. We then compared the spectra obtained for the proteins alone and in presence of amylose and amylopectin. The CD spectra for both proteins alone have been described in (Liu, Pfister, Osman, Ritter, Heutinck, Sharma, Eicke, Fischer-Stettler, et al. 2023) and show that both proteins are structured with differences in the secondary structure composition. For ESV1, the pattern of CD spectra corresponds to a folded protein with a strong positive band at λ=196nm and only one negative band at λ=220nm, which is characteristic of all β-proteins. For LESV, the pattern of the spectrum reveals a global folding of β-strands and α-helices. Indeed, the LESV CD spectrum shows a strong maximum at λ=192nm and a minimum at λ=216nm, which are the signature of the presence of β- structures, but unlike ESV1, the spectrum also shows two shoulders at λ=210 and λ=222nm, which are evidences for the concomitant presence of α-helices. These results are consistent with models generated by Alphafold that predict the folding of both proteins into a conserved β -sheet C-terminal domain and the presence of some α-helices in the N-terminal part of LESV (Liu, Pfister, Osman, Ritter, Heutinck, Sharma, Eicke, Fischer-Stettler, et al. 2023).

To analyze the interactions between proteins (ESV1 and LESV) and polyglucans (amylose and amylopectin), 4 µl of solutions of each protein were mixed with 1 µl of 1% amylose or 1% amylopectin solutions and incubated for 2 h before measurement. For each spectrum, the composition of the secondary structure elements was determined using BestSel (Micsonai et al. 2015). The values obtained were compared with those obtained for the proteins alone.

**Figure 3** shows the superposition of the spectra of LESV alone and added to amylopectin or amylose solutions. The spectrum obtained for the LESV/amylose mixture has a broadly similar appearance to that of LESV alone, with peaks of slightly lower magnitude, showing that the presence of amylose induces no or only subtle conformational changes in the protein.

**Figure 3:**
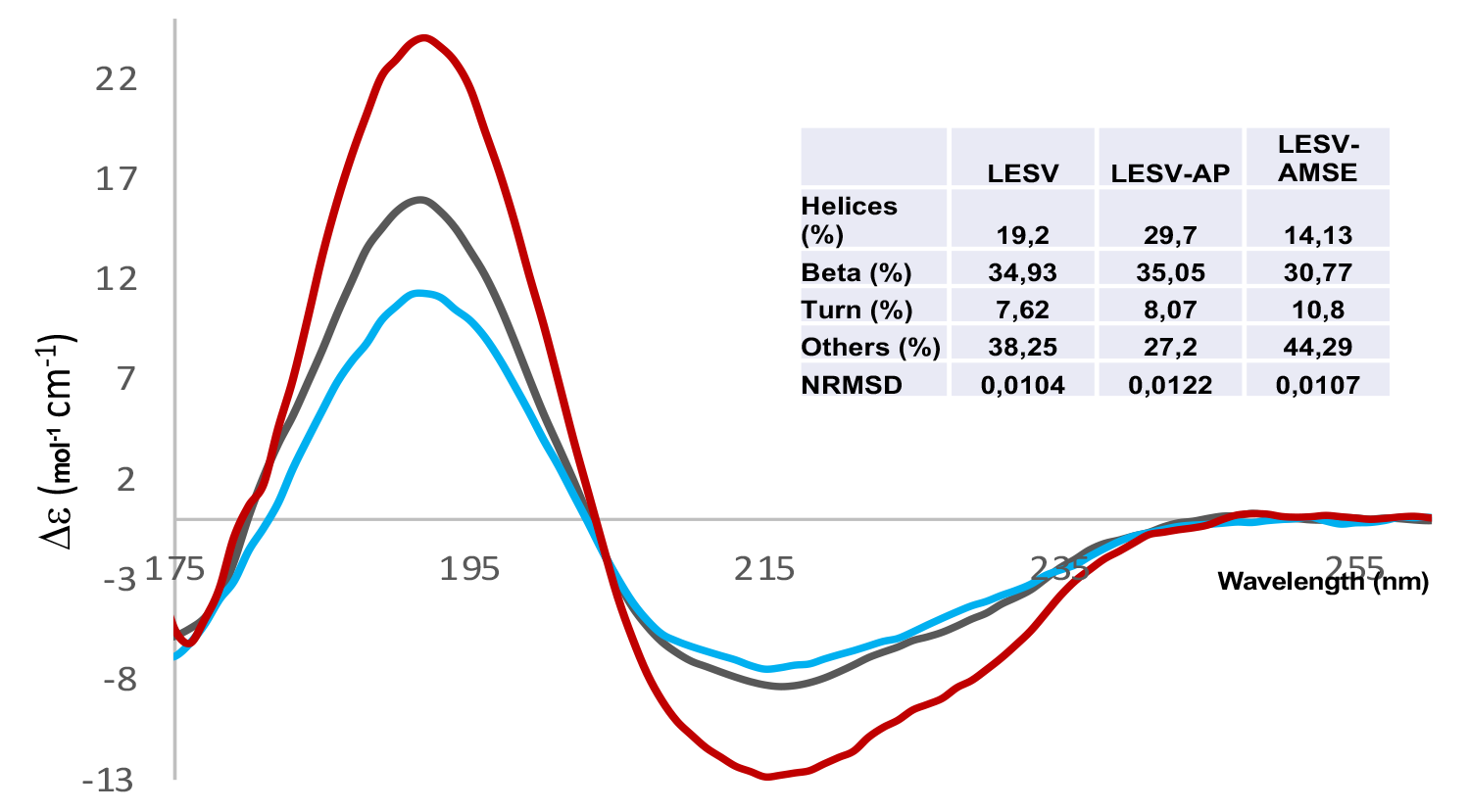
*SR-CD spectra for LESV* alone (grey), and in presence of amylopectin (red) or amylose (blue). The composition in secondary structural elements evaluated by BestSel are in inset.

In contrast, the spectrum obtained for the LESV/amylopectin mixture shows peaks of much higher magnitude than those obtained for the protein alone or in the presence of amylose. The positive peak at λ=192 nm has a magnitude 60% higher than that of the protein alone, indicating a more significant structuring of the protein. At λ=208 and λ=215 nm, the molar ellipticity values indicating the presence of α and β structures, are 50% lower than those observed for the protein alone or in the presence of amylose. More interestingly, the molar ellipticity at λ=222 nm, indicating the presence of α-helices, is much lower (70%) than that observed for the protein alone or in the presence of amylose. This result shows that the binding of amylopectin induces a significant conformational change of LESV notably through the formation of additional α-helices upon binding of amylopectin. On the other hand, the presence of amylose does not induce any structural modification, confirming EMSA experiments showing no affinity for this glucan.

To identify and quantify the conformational changes undergone by LESV in the presence of amylose and amylopectin, the composition of the secondary structural elements was analyzed using BestSel (Micsonai et al. 2015). The results presented in inset in **Figure 3** confirm the analysis of the CD spectra. The composition of the β-strands and turns is equivalent whether LESV is alone or in the presence of amylose or amylopectin, indicating that the structure of the β-domain is not modified by the presence of polyglucans. LESV in the presence of amylose seems to have a slightly lower number of α-helices and strands than the protein alone. On the other hand, a higher number of α-helices is observed when the protein is in the presence of amylopectin. This result confirms that the LESV protein interacts with amylopectin and that this interaction induces the formation of α-helices, most probably located in the N-terminal domain, as no changes in the content of β-strands attest to the preservation of the structure of the C-terminal domain.

The same analysis was performed for ESV1. The analysis shows broadly equivalent spectra for ESV1 in the presence of amylose or amylopectin, with a degree of structuring that appears to be slightly lower for the protein alone. Analysis of the composition of secondary structural elements by BestSel, displayed in inset in Figure 4 indicates an equivalent composition for ESV1 alone or in the presence of amylose and a slight reduction or modification of some strands and α-helices when the protein is in the presence of amylopectin. This result indicates that ESV1 interacts with amylopectin without inducing significant conformational change in the protein.

**Figure 4:**
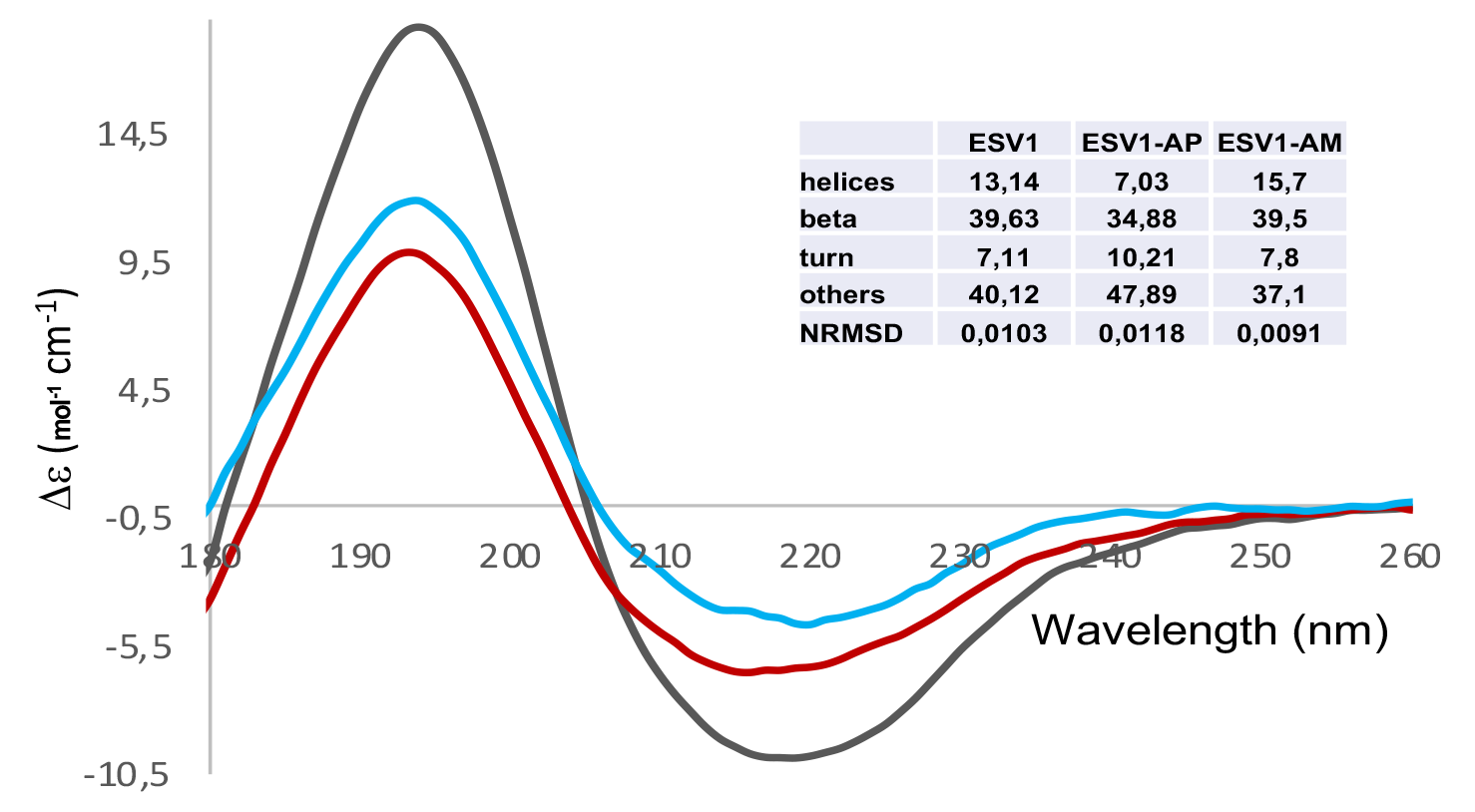
*SR-CD spectra for ESV1* alone (grey), and in presence of amylopectin (red) or amylose (blue). The composition in secondary structural elements evaluated by BestSel are in inset.

### Glucan binding affects the Melting temperature (TM) of LESV but not of ESV1

Enhanced detection of ligand binding can be achieved through thermal denaturation studies monitored by SR-CD. This method is more sensitive than simple spectral differences as it can detect interactions that do not induce structural modifications of the proteins. By employing this method, it is possible to determine the melting temperature (TM) of the unfolding transition. The CD signal variation was measured as a function of the temperature for both proteins, alone and in the presence of amylose or amylopectin, under the same conditions as for the spectra at constant temperature.

Figure S2 presents the results obtained for LESV. During protein denaturation, two stable states were observed, either alone (figure S2A) or in the presence of amylopectin (figure S2B) or amylose (Figure S2C), which were expressed by an isosbestic point at λ=210 nm. As temperature increased, the intensity of the positive peak at λ=192 nm decreased, and a hypsochromic effect (shift towards lower wavelengths) was observed, reflecting significant modifications of the global protein structure. The magnitude of the peak between λ=210 and λ=225 nm decreased sharply, indicating a decrease in the α and β structures. Particularly the decrease at λ=222 nm reflected a greater decrease in the α-helix structures in the LESV amylopectin mixture. Simultaneously, a new negative band appeared at λ=203nm, indicating the formation of random coils and thus attesting to the denaturation of the protein.

To assess the impact of the presence of glucans on protein stability, we measured the TM of the mixtures by monitoring the molar ellipticity evolution as a function of temperature at λ=195nm. The curve obtained have been normalized and are presented in figure 5A. The experiment shows that the denaturation of LESV is slowed down in the presence of amylose and even more so in the presence of amylopectin with TMs of around 55°, 60° and 65° respectively. This result shows that the presence of amylose and amylopectin stabilizes LESV, suggesting an interaction of the protein with the two starch components.

**Figure 5:**
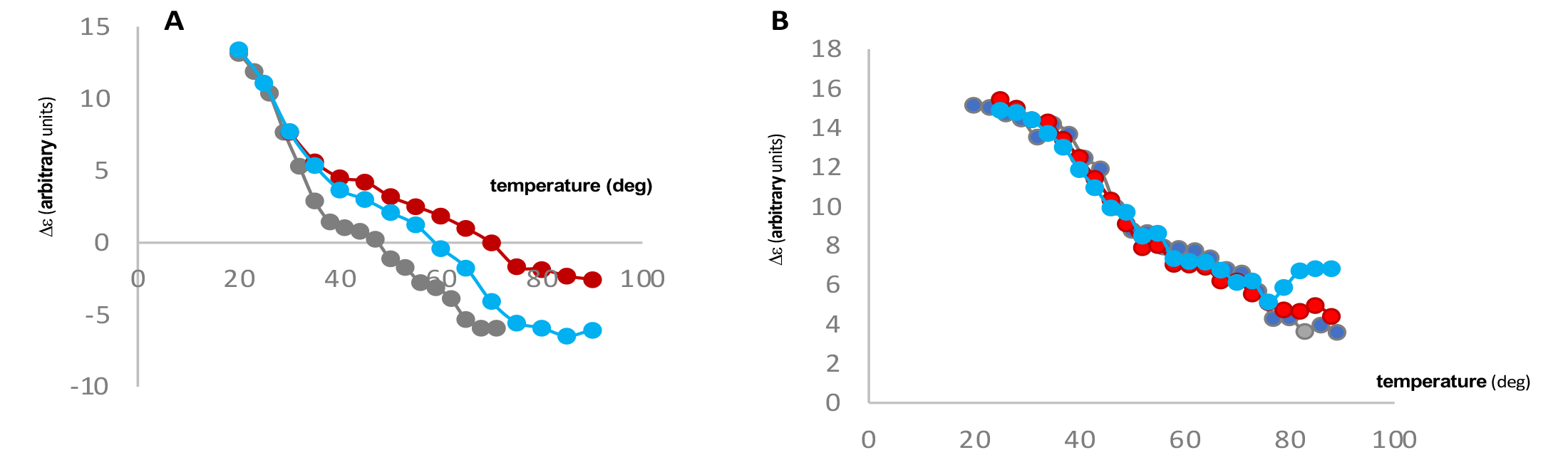
*Thermal denaturation of LESV and ESV1 followed by the variation of the molar ellipticity in function of the temperature.* A) thermal denaturation of LESV followed at l =190 nm B) thermal denaturation of ESV1followed at l=195nm. Curves corresponding to proteins alone, with amylopectin or amylose are colored in grey, red and cyan respectively.

The experiment was replicated with ESV1, and the results were analysed in figure S2. It was observed that two isosbestic points were present for the protein alone (Figure S2D) with a first isosbestic point at λ=210nm as observed for LESV and a second one at λ=197nm. The same profile was observed for the protein in the presence of amylopectin (Figure S2E). This observation confirms that denaturation of the protein alone or in the presence of amylopectin occurs in two different states. Only one state was observed in the presence of amylose (Figure S2) where only the isosbestic point at λ=201nm was present suggesting that the presence of amylose may stabilizes ESV1 in the first step of denaturation.

In all assays, as a result of the increasing temperature, the peak observed at λ=193 nm decreases in intensity and undergoes a hypsochromic effect before stabilizing around λ=180nm. After the first step of denaturation this peak continues to decrease in intensity without undergoing a hypsochromic effect, indicating a change in the global protein structure. The intensity of the negative peak observed at λ=220 nm decreases and also undergoes a hypsochromic effect (that stabilize around λ= 212nm) until the first denaturation state obtained at 43°, 43° and 46° degrees for ESV1, ESV1/amylopectin and ESV1/amylose respectively. At higher temperatures a different behavior is observed for the ESV1/amylose mixture. The size of the negative peak stops to decrease, while the appearance of a negative peak around λ=202 nm is observed, a peak that is not observed for ESV1 alone or in complex with amylopectin. For ESV1 alone and in complex with amylopectin, a second state is reached at about 75°. Between the first and second states, the positive peak at λ=180nm continues to decrease in intensity without undergoing a hypsochromic effect. Similarly, the peak at λ=212nm continues to decrease in size without shifting.

Looking at the molar ellipticity evolution as a function of temperature at λ=190nm. The curve obtained have been normalized and are presented in figure 5B. The denaturation curves show the same pattern and can be overlaid with an inferred TM of about 50°C for ESV1. Thus, in contrast to LESV, the presence of amylose or amylopectin does not affect the thermostability of ESV1, despite a demonstrated interaction with amylopectin.

### ESV1 and LESV accumulate on the entire surface of starch granules

To confirm the results obtained in solution, the binding of ESV1 and LESV to starch granules was visualized by UV fluorescence microscopy. This approach, which allows proteins to be visualized by the fluorescence of their aromatic residues on starch granules without adding any external probe, has been used and described in (Tawil et al. 2011). Starch granules contain a large amount of GBSS, the enzyme responsible for amylose synthesis. To carry out the experiment, we used starch granules from maize in which the GBSS gene was deleted, so that the fluorescence of the granules could be distinguished from the fluorescence of the protein being studied (see methods part). The measurement was carried out simultaneously in visible light and with excitation at λ=310 nm, which allows the tryptophans residues to be exited. The emission spectrum was obtained using a filter to select a wavelength range between 329 and 351 nm. Two controls were carried out, the first with starch granules alone to verify the absence of fluorescence and the second with starch granules in the presence of BSA to verify the absence of unspecific protein binding. 2µl of WAXY starch grains (300 mg/ml) were incubated with 2µl of ESV1 (4mg/ml) or 1µl of LESV (10mg/ml) for 2 hours before measurement. Visible and fluorescence images are shown in Figure 6. The fluorescence images clearly show a distinct halo over the entire surface of the starch granules. This fluorescence corresponds to the emission at the selected wavelength of the tryptophan residues present in ESV1 and LESV. These images demonstrate that the proteins accumulate on the surface of the starch granules with no difference in location between ESV1 and LESV and confirm the affinity of both proteins for amylopectin in its insoluble form.

**Figure 6:**
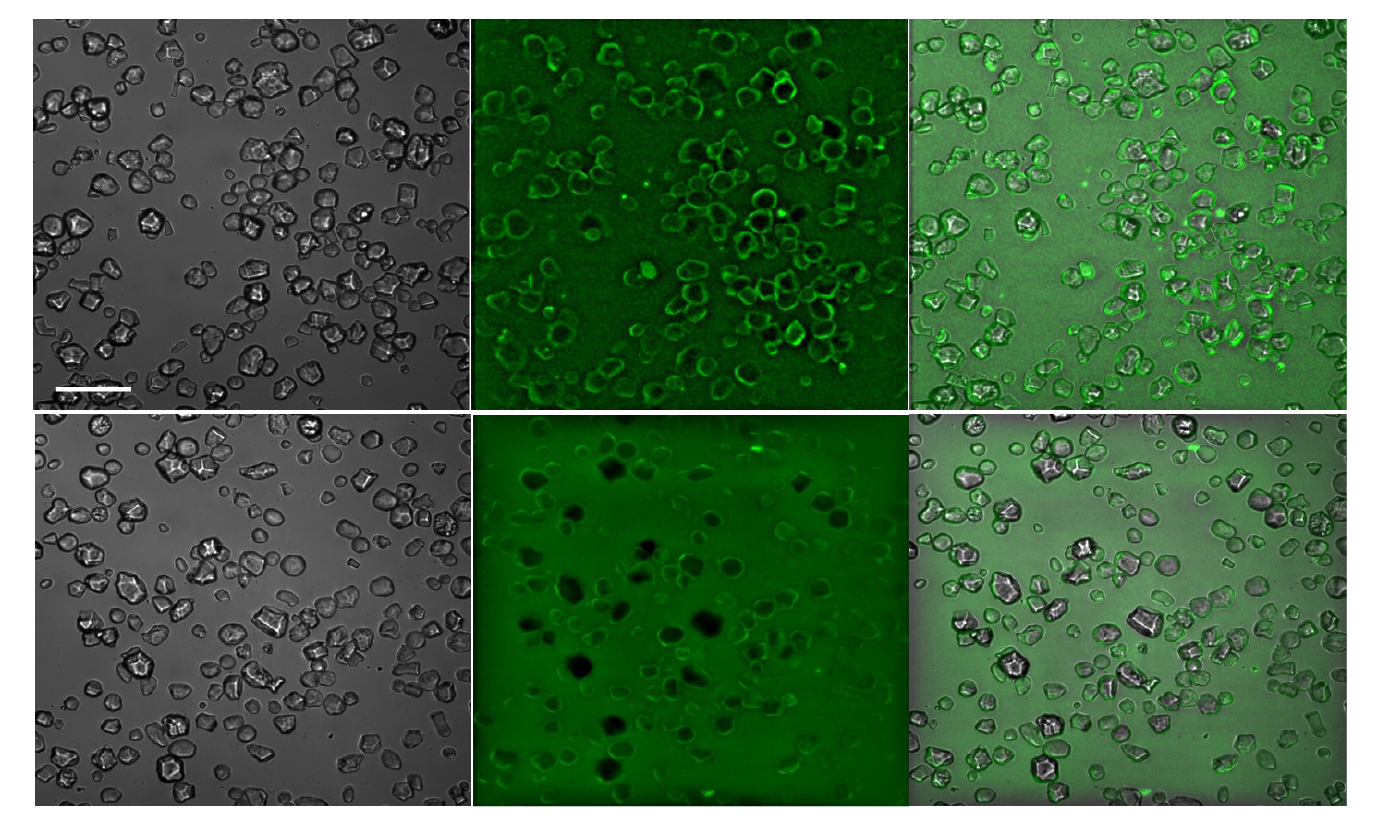
Visible light and fluorescence imaging of maize waxy starch granules in presence of ESV1 (top panel) or LESV (bottom panel). A) visible light imaging B) fluorescence images of ESV1 or LESV absorption on starch granules C) combination of visible light and fluorescence images.

### The C-terminal tryptophan rich domain of ESV1 and LESV can bind two double helices of amylopectin (at least) on one face of the β-sheet

To gain more insight into the interaction of the new sugar binding domain (SBD) identified on ESV1 and LESV, we simulated the complex with the conserved C-terminal domain of the proteins and two double helices of amylopectin. To do this, we first used the model of LESV containing the β-sheet and the conserved and well positioned α-helix described in (Liu, Pfister, Osman, Ritter, Heutinck, Sharma, Eicke, Fischer-Stettler, et al. 2023) and shown in Figure 1. In models of both proteins, we identified a polarity on both sides of the C-terminal β-sheet, one of which is occupied by a long helix (LESV) or a long loop (ESV1) that partially obscures the aromatic or acidic amino acid residues that could interact with amylopectin (Supplementary fig S1). In this work, we have studied the interaction of amylopectin only on the accessible side of the β-sheet. The N-terminal domain, for which no structure has been predicted was omitted. We first performed a docking calculation with one molecule of protein and one double helix of amylopectin centred on one aromatic stripe of LESV on the accessible face of the β-sheet. We obtained a good solution in which one double helix of amylopectin binds the β-sheet which is well aligned with the aromatic stripe. We repeated the same approach with the protein binding a single chain of amylopectin as the target centred on the second aromatic stripe. We again obtained a solution shown in figure 7. On this structure, two double helices of amylopectin bind along the aromatic stripes and lie parallel to each other separated with 10 Å (between the axes of the double helices). This arrangement of the double helices in relation to each other corresponds to the arrangement of the amylopectin molecules described for the type A-type starch which can be mostly found in plant leaves (Imberty et al. 1988). This result demonstrates that the conserved C-terminal domain conserved of ESV1 and LESV is perfectly suited to the interaction of the proteins with the semi-crystalline form of native amylopectin.

**Figure 7:**
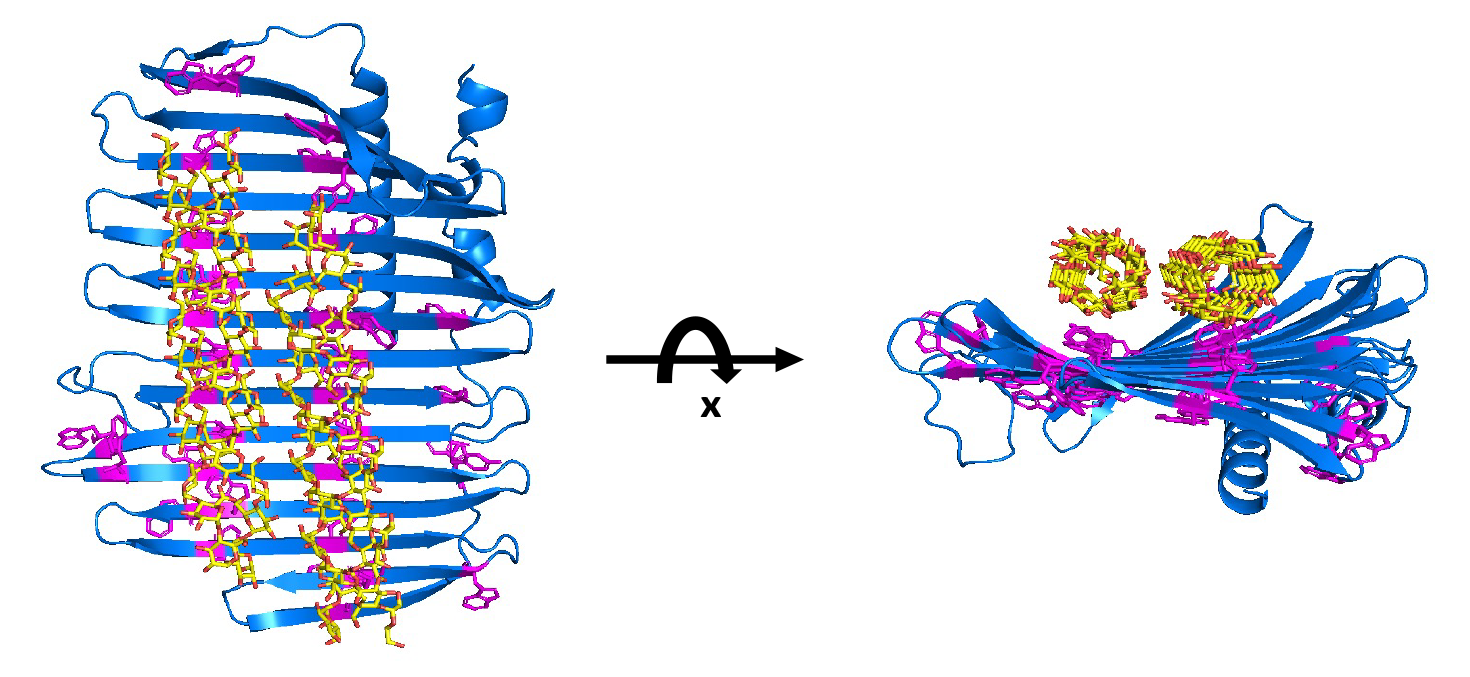
Molecular model of the complex between C-terminal domain of LESV and amylopectin double helices. Protein chain is represented in cartoon and colored in blue. Aromatic residues are colored in magenta and their side chains are represented by sticks. Amylopectin double helices are represented by sticks and colored by atom types.

**Figure 7:**
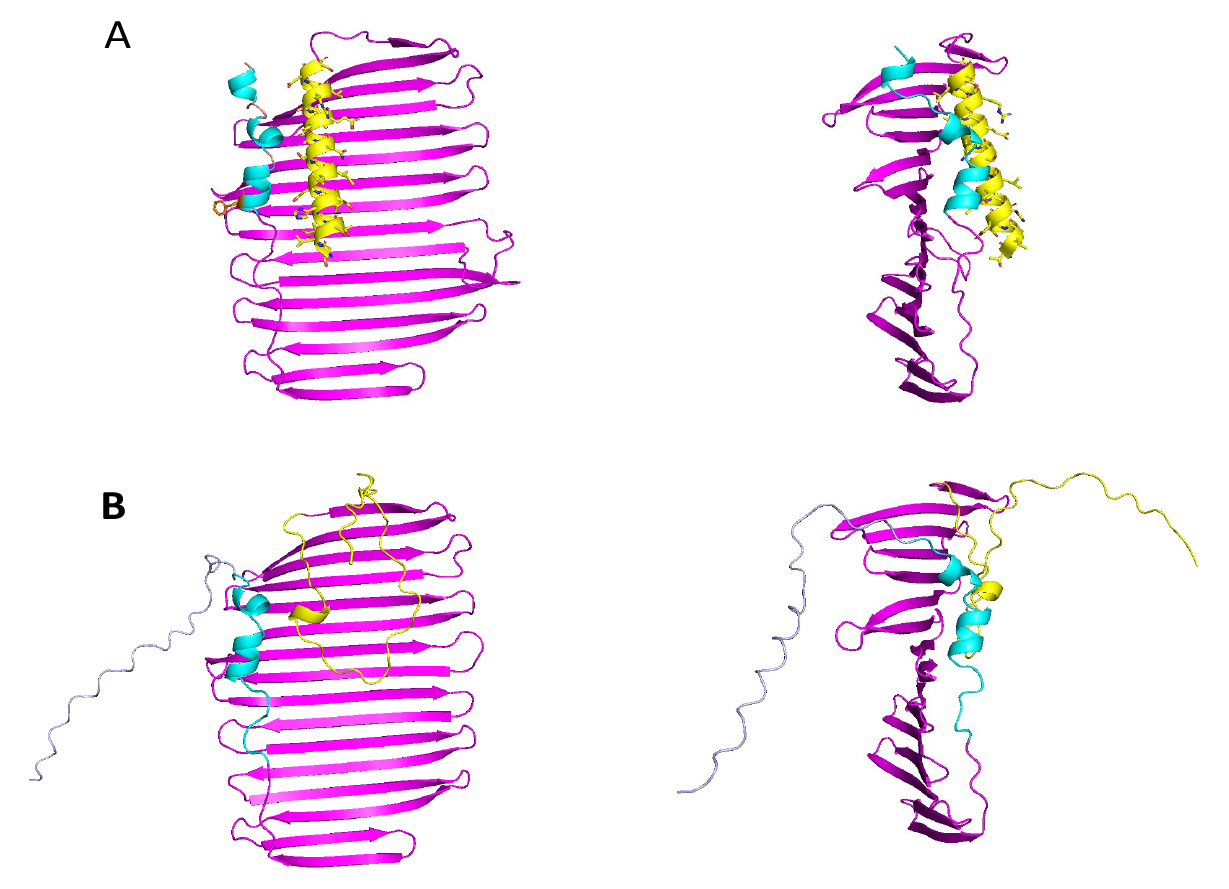
Molecular model of the complex between C-terminal domain of LESV and amylopectin double helices. Protein chain is represented in cartoon and colored in blue. Aromatic residues are colored in magenta and their side chains are represented by sticks. Amylopectin double helices are represented by sticks and colored by atom types.

The amylopectin molecules interact with the protein domain through numerous interactions typical of protein-sugar interactions, as predicted from the analysis of the primary sequence of the β-domain and the distribution of conserved amino acids in the Alphafold model. The glucose rings can interact by hydrophobic stacking with the aromatic rings of the β-domain all along the double helices. The acidic residues conserved in LESV and ESV1 also play an important role in the interaction by participating in hydrogen bonding with the hydroxyl groups of the glucose residues on the sides of the double helices. On face A, described above, there is an analogous organization in stripes of aromatic and acidic residues, suggesting that double helices could also bind on this face after structural reorganization of the conserved helix (LESV) or loop (ESV1) allowing the proteins to insert between amylopectin molecules.

## Discussion

### ESV1 and LESV bind specifically to amylopectin

In this work, we have investigated the interaction specificities of ESV1 and LESV with the different components of starch. The specificities of amylopectin and amylose macromolecules, which are complex, non-homogeneous and often dense glucans in solution, limited the approaches that could be used. We first chose an EMSA approach, which allowed us to analyze the behavior of both proteins in relation to each of the polyglucans. Previous work on the affinity of ESV1 and LESV for starch components (Malinova et al. 2018; Singh et al. 2022) focused on the interaction between ESV1 and LESV and starch glucans in insoluble form and/or on different mutant starch granules. This work proposed that ESV1 and LESV interact with starch granules, each having specific affinity for amylopectin or amylose respectively, this affinity being independent on the protein/glucan ratio. These results are, however, not fully consistent with the recent characterization of LESV and ESV1 (Liu, Pfister, Osman, Ritter, Heutinck, Sharma, Eicke, Fischer-Stettler, et al. 2023) and with our results. To remove this ambiguity, we further analyzed the interaction between ESV1 and LESV, taking into account different parameters. By studying the migration profile of the two proteins in the presence of different glucans, we were able to show that both proteins have a strong affinity for amylopectin and that this affinity increases with the amount of this glucan. We also didn’t find any differences in the behavior of the two proteins towards amylose. Indeed, in the presence of amylose, we did not observe any migration delay associated with amylose or other linear polyglucans using this approach. Structural analysis of ESV1 and LESV in the presence of amylose and amylopectin by SR-CD shows that only the presence of amylopectin induces a significant conformational change in LESV upon interaction, leading to the structuring of disordered regions of the N-terminal domain of the protein into α- helices. No conformational changes were observed in ESV1, but this can be explained by the fact that the protein has a reduced N-terminal domain and consists almost entirely of the highly structured C-terminal conserved domain, which is unlikely to undergo conformational changes. However, we observed that the presence of amylose increased the TM of LESV although to a lesser extent than amylopectin. All these observations suggest that amylose is able to interact with LESV and ESV1, but that this interaction, unlike that with amylopectin, is not very specific and is probably related to the abundance of aromatic amino acids in the structure of both proteins.

As the experiments we carried out were with amylopectin and amylose molecules in their solubilized form (obtained by chemical treatment), we wanted to verify that ESV1 and LESV were able to interact with amylopectin in its crystallized form. The results we obtained in fluorescence microscopy with maize *waxy* starch granules, which contain no amylose, clearly showed that ESV1 and LESV interact directly with amylopectin starch granules. In fact, the two proteins, identified by their fluorescence, accumulate on the entire surface of the starch granules. This result confirms the results obtained in solution and is consistent with the role described in our recent work.

### LESV is able to bind to amylopectin during its biosynthesis

The two proteins behave differently toward glycogen and only LESV was able to interact with it. This finding is interesting in several levels. First, the difference in affinity of ESV1 for amylopectin and amylose does not seem to be related to the presence of branching points, but rather to the three-dimensional structure of the glucan double helices. Secondly, an analogy have been drawn between the structure of glycogen and that of amylopectin during biosynthesis, before the action of isoamylases that remove the excess branching points that are not compatible with the formation of the final cluster-structure of amylopectin(Ball et al. 1996). This result confirms the proposed function of LESV in our previous work (Liu, Pfister, Osman, Ritter, Heutinck, Sharma, Eicke, Fisher-Stettler, et al. 2023), where LESV would intervene directly upstream (or concomitantly) of the action of isoamylases during the biosynthesis of amylopectin chains. LESV would be able to support the phase transition of double helices, although their number of branch points has not been optimized. The fact that LESV has an affinity for glycogen, which has a similar structure to amylopectin chains before the action of isoamylases, is consistent with the proposed function of LESV. This result is also consistent with the proposed function of ESV1, which would act downstream of LESV to stabilize the newformed starch granules and has an affinity only for the amylopectin chains in the crystalline phase of the starch granules.

LESV and ESV1 share a common domain whose structure was described as being particularly compatible with the binding of amylopectin double helices. The fact that ESV1, which is predominantly composed of this domain, does not interact with glycogen suggests that the interaction with the forming amylopectin is mediated by the N-terminal domain of LESV, which gives LESV additional specificity by allowing it to accommodate different glucans than ESV1.

### C-terminal domain is designed to bind specifically (at least) two double helices of amylopectin

The models obtained for ESV1 and LESV *via* Alphafold provided a very reliable structure for the C-terminal tryptophan-rich domain, which is conserved between the two molecules. The other parts of the molecule were predicted without much reliability (Liu, Pfister, Osman, Ritter, Heutinck, Sharma, Eicke, Fischer-Stettler, et al. 2023). The C-terminal domain folded into an original structure, forming a rather large oval (about 40 Å wide and 70Å long) antiparallel twisted ß-sheet. On this β-sheet, the aromatic and acidic residues, organized in repeated sequences identified during the analysis of the protein sequences, form parallel lines equidistant from each other and parallel to the axis of the β-sheet. The side chains of these amino acids point alternately to both sides of the ß-sheet, suggesting a bipolarity of the ß- sheet with respect to the attachment of amylopectin chains. However, one of the faces of the β-sheet is occupied by a conserved-C-terminal helix on its side and a long α-helix or a long loop in LESV and ESV1 respectively. Based on our current data, even though we have observed protein conformational changes in the presence of amylopectin, we do not know if these parts can move to release the aromatic and acidic residue lines for amylopectin molecule binding. If that was the case, and if ESV1 and LESV could bind amylopectin on both sides of their β-sheet, we would expect to find assemblies with sandwich-like alignments of proteins and amylopectin. However, we have never observed such structures during the interaction experiments. Therefore, we computed models where only the "free" side interacted with double helices of amylopectin. We were able to demonstrate that LESV and ESV1 can bind amylopectin in both its soluble form and its organized form within starch granules. Furthermore, we have shown that this interaction occurs through the common domain shared by LESV and ESV1, which is organized in a β-sheet structure using conserved aromatic and acidic residues. Moreover, we have shown that the unique and unprecedented structure of this domain enables it to bind two parallel double helices of amylopectin in the arrangement found in the crystalline phases of starch granules, thereby enabling its function in the organization and maintenance of starch granules in plants.

### The N-terminal domain allows regulation of LESV specificity towards starch components

The structure of the N-terminal domain of LESV is not known. However, we have been able to demonstrate that it consists mainly of disordered regions and α-helices organized close to the C-terminal domain. We have also shown that interaction with amylopectin induces the formation of α-helices from disordered regions without affecting the β-sheet structure. In ESV1, this domain is highly reduced but still predicted to be unstructured, as well as the polyproline tail at the C-terminus. It is likely that the presence of these regions prevented us from obtaining crystals. Since amylopectin structure and size is not monodisperse in solution, it cannot be used to stabilize the proteins, and we are currently searching for analogues that can facilitate the crystallization of both proteins. While analyzing the structure of the C-terminal domains has allowed us to describe the interaction mode of the two proteins with amylopectin, analyzing the entire structures, particularly for LESV, would enable us to precisely describe the function of the N-terminal domain, even though we have shown that it is likely involved in the difference of specificity of both proteins. Considering that several helices are present in this domain in LESV and taking into account its involvement in amylopectin biosynthesis, it is possible to consider that LESV may also interact with other proteins, as already described for SSs and BEs involved in starch biosynthesis (Crofts et al. 2015; Ahmed et al. 2015).

This work will enable us to improve our understanding of the molecular mechanisms of ESV1 and LESV in Arabidopsis. We have shown that the C-terminal domain, which is conserved in both ESV1 and LESV, is particularly well suited to binding amylopectin double helices as they are organized in starch granules. We have also shown that LESV is able to interact with amylopectin molecules during biosynthesis. These results suggest that its involvement in the phase transition probably occurs before the end of double helix biosynthesis. ESV1 would intervene after the biosynthetic process to stabilize the granules and prevent premature degradation by degradative enzymes. This work also showed that the N-terminal domain of LESV undergoes folding induced by interaction with amylopectin, including the structuring of unfolded regions into α-helices, and that it is likely to be involved in regulating the function of the protein. Further research is needed to describe the precise molecular mechanism of ESV1 and LESV in plants. It is clear that the resolution of the atomic structures of ESV1 and LESV interacting with starch glucans will allow considerable progress to be made in this area, as well as in studying the function of these proteins in other organisms and their involvement in storage starch biosynthesis.

## Aknowledgements

The authors strongly acknowledge DISCO beamline and the regular access to the small angle X-ray scattering beamline SWING at synchrotron SOLEIL (St Aubin, France) through the BAG MX-20181002 and MX-20201190, and are grateful for the expert technical support provided by beamline staffs.

## Author Contributions

CB and RO conceived and designed the experiment with imput from SZ, DD and CdH. RO and CB performed experiments with imput from MB, CL and CS. CB, RO and MB collected synchrotron data. CB and RO analyzed data with the imput of MB and DD. CB wrote the manuscript. CB, DD, CdH, SZ and CS revised the manuscript.

## Supplemental figures

**Figure S1:** Structural conserved motifs on the Face A of LESV (top) and ESV1 (bottom). The structures are represented as cartoon, the common β-sheet is colored in magenta, the common C-terminal helix is colored in cyan and the long helix of LESV and the long loop of ESV1 are colored in yellow. The right panel represents the left panel after a rotation of 90° along y-axis.

**Figure S2:**
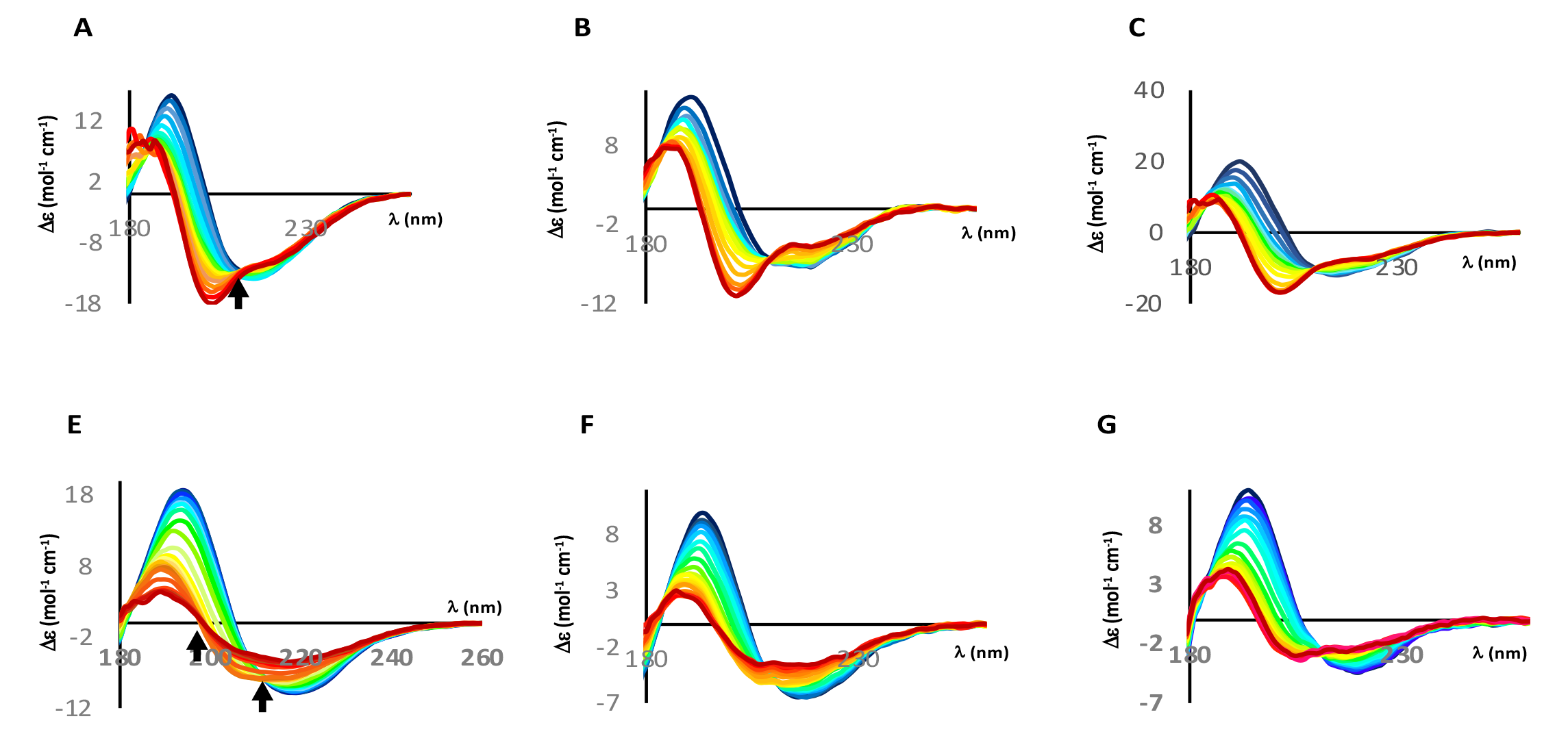
Thermal denaturation of LESV and ESV1 followed by SR-CD. Each plot represents consecutive scans on the protein collected at a set of temperature between 20 to 90°. Scans are colored in a gradient from dark blue (first temperature) to dark red (last temperature) for A) LESV alone, B) LESV with amylopectin C) LESV with amylose, D) ESV1 alone, E) ESV1 with amylopectin and F) ESV1 with amylose. Black arrows indicate the position of isosbestic points.

